# Proteome profiling reveals HES1-driven mitotic catastrophe in ovarian serous carcinoma

**DOI:** 10.1101/2025.08.20.671256

**Authors:** Jie Bao, Sanna Pikkusaari, Jun Dai, Samuel Leppiniemi, Wenjun Huang, Weiming Yang, Anu Anil, Mirva Pääkkönen, Chuqi Lei, Eva Daniela Mendoza-Ortiz, Ezgi Karagöz, Johanna Eriksson, Min Li, Johanna Hynninen, Otto Kauko, Anniina Färkkilä, Anna Vähärautio, Sampsa Hautaniemi, Liisa Kauppi, Jing Tang

## Abstract

Ovarian high-grade serous cancer (HGSC) is an aggressive subtype of epithelial ovarian cancer. Here, we identify BX-912, a phosphoinositide-dependent kinase 1 (PDPK1) inhibitor, as a promising therapeutic agent for HGSC. BX-912 suppressed HGSC growth as a single agent and synergized with olaparib independently of BRCA status. Unexpectedly, BX-912 treatment induced multinucleation, a phenotype not observed with other PDPK1 inhibitors. Proteome Integral Solubility Alteration (PISA) profiling revealed the transcription factor HES1 as a functional target of BX-912. Structural modeling showed that BX-912 binds the Orange domain of HES1, while its WRPW motif mediates interactions with protein partners, including the AP2 endocytic protein complex, coordinating their nuclear accumulation that leads to a mitotic catastrophe. Furthermore, cell cycle analyses showed that BX-912 combined with olaparib synergistically enhanced DNA damage and G2-M arrest. Our study demonstrates the value of proteomics for revealing hidden drug activities. It also identifies potential inhibition strategies for HES1, which is commonly overexpressed in HGSC. Additionally, this study proposes a novel strategy of targeting consecutive cell cycle phases to enhance treatment efficacy in HGSC.

## Introduction

It is well-established that drugs can bind to multiple proteins, potentially causing unwanted side effects [1] or off-target effects [2]. Systematic mapping of drug-target interactions can be achieved via various *in silico* or *in vitro* techniques [1, 3–5]. Experimental target identification methods range from conventional protein pull-down experiments to advanced approaches that monitor changes in cellular, chemical, or structural properties following drug treatment [6]. Over the past decade, proteomics assays using thermal protein profiling (TPP) have been developed for large-scale drug target deconvolution in live cells [7]. Based on the principle central to the cellular thermal shift assay [8], TPP identifies drug-binding proteins by leveraging mass spectrometry to detect changes in protein solubility across samples heated to incremental temperatures, reflecting shifts in aggregation temperature due to drug binding. A recent advancement of TPP, known as the proteome integral solubility alteration (PISA) assay, enables the pooling of soluble proteins obtained at different temperatures in a single run. This improvement increases the throughput of target identification by 10-fold, providing higher statistical power with reduced sample amounts [9–11]. However, as a relatively new method, questions remain regarding whether this increased scalability compromises accuracy, and how effectively the proteomics profiles can lead to novel findings in drug repurposing and clinical translation.

Kinase inhibitors frequently exhibit polypharmacological or off-target effects due to their primary interaction with conserved ATP-binding pockets [12]. An illustrative example is the PDZ-binding (for anchoring membrane-bound receptors to cytoskeletal components) kinase inhibitor OTS964, which was found to instead target cyclin-dependent kinase CDK11, as demonstrated by a CRISPR/Cas9-mediated genetic essentiality assay [13]. Kinome profiling has also revealed that different generations of PARP inhibitors—a class of drugs that have transformed the treatment of HR-deficient malignancies, including ovarian high-grade serous cancer (HGSC)—exhibit distinct polypharmacological profiles [14, 15]. Nonetheless, there remains a significant gap in understanding how secondary effects of kinase inhibitors may arise through interactions with non-enzymatic proteins, such as transcription factors and scaffold proteins. Addressing this challenge may necessitate real-time capture of protein interactions, which can be optimally accomplished through proteomics-based target profiling [16].

3-phosphoinositide-dependent protein kinase-1 (PDPK1) is a master kinase orchestrating multiple oncogenic pathways, including AKT, MAPK, and MYC [17]. Overexpression of PDPK1 has been documented in various malignancies and is notably implicated in regulating chemoresistance in ovarian cancer [18]. Multiple PDPK1 inhibitors have been developed either as monotherapy or as part of combinatorial therapeutic regimens [19]. Despite the theoretical appeal of targeting such a key oncogenic kinase for cancer treatment, no PDPK1 inhibitors have been applied in clinical settings.

In this study, we identified BX-912, a putative PDPK1 inhibitor, either alone or in combination with the PARP1 inhibitor olaparib, effectively induced cytotoxicity and proliferation arrest in HGSC cell lines. Notably, this efficacy was not observed in other PDPK1 inhibitors. Proteome Integral Solubility Alteration (PISA) analysis and *in vitro* validation experiments identified the transcription factor HES1 as the functional target of BX-912 in HGSC, mediating cell cycle arrest. These findings underscore the importance of elucidating drug targets to understand their mechanisms of action and highlight the potential of targeting HES1, in a non-canonical manner, to induce mitotic catastrophe in HGSC treatment.

## Results

### BX-912 demonstrates a uniquely potent cytotoxic effect on HGSC cells

PDPK1 plays an essential role in regulating oncogenic signaling pathways, including AKT, MAPK, and Myc [17] (Fig.1A). To explore its therapeutic potential, we conducted a high-content drug screen using three HGSC cell lines, including olaparib-sensitive (COV362), intrinsically resistant (Kuramochi), and acquired-resistant (COV362 re-expressing *BRCA1*: COV362+B1). We evaluated three PDPK1 inhibitors with known anti-tumor activities (Fig.1A). Among them, BX-912 [38] exhibited significant cytotoxic effects in a five-day treatment. Unlike the PARP inhibitor olaparib, the efficacy of BX-912 was not influenced by cell homologues repair (HR) status, with an IC_50_ of approximately 0.5 µM observed across all three tested HGSC cell lines (Fig.1B). Additionally, a synergistic effect between BX-912 and olaparib was observed (Fig.1C, Fig EV.1A). These findings were further validated in a long-term three-week colony formation assay (Fig.1C, Fig EV.1A).

**Figure 1.**
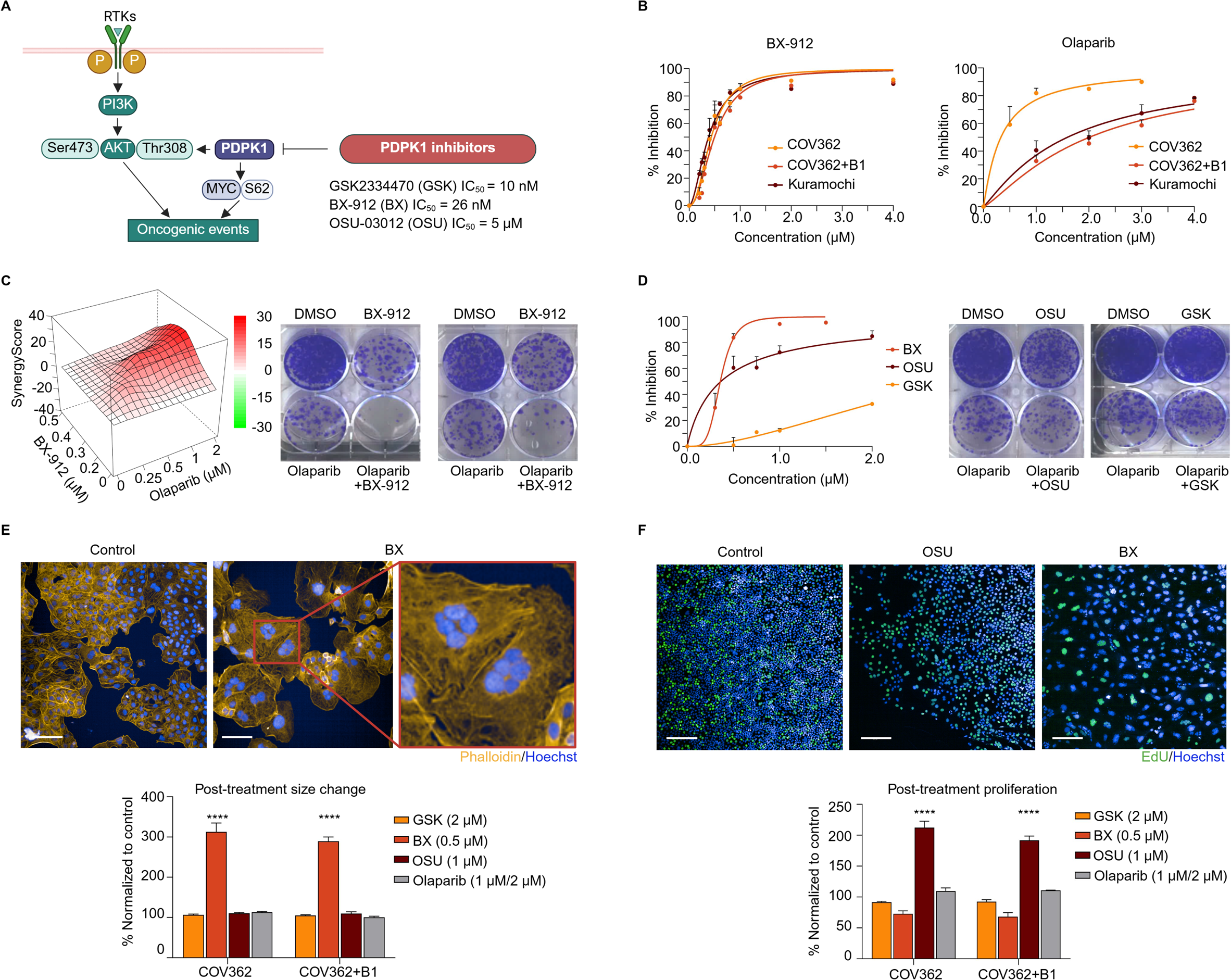
BX-912 exhibits a distinct cytotoxic effect on HGSC cells compared to other PDPK1 inhibitors. A Schematic representation of the PDPK1 pathway and its downstream targets, along with the three PDPK1 inhibitors (BX-912, OSU-03012, and GSK2334470). The IC_50_ values represent the concentration at which each drug achieves 50% inhibition of PDPK1 kinase activity in cell-free assays. B Dose-response curves for BX-912 and olaparib in three HGSC cell lines following a five-day treatment. C Left: Drug synergy scores of BX-912 and olaparib in olaparib-resistant COV362 cells re-expressing BRCA1 (COV362+B1). Mean HSA score: 10.71, P = 3.2x10^35^. Right: Long-term recovery and colony formation assay demonstrating the synergistic effect of BX-912 (0.4 µM) and olaparib (1.0 µM) treatment. D Left: Dose-response curves of the three PDPK1 inhibitors in COV362+B1 cells. Right: Long-term colony formation assay showing the lack of efficacy of OSU-03012 (0.5 µM) and GSK2334470 (1.0 µM) in COV362+B1 cells. Olaparib was used at 1.0 µM. E Representative images and corresponding nuclear morphological changes following the drug treatments. COV362 cells were treated with 1.0 μM olaparib while COV362+B1 cells were treated with 2.0 μM olaparib due to their differential sensitivity to the drug. **** P < 0.0001, one-way ANOVA analysis comparing BX-912 treatment to others. Blue: nucleus; yellow: cytoskeleton. Scale bar: 100 µm. F Representative images and bar plot depicting proliferation rates following different drug treatments. **** P < 0.0001, one-way ANOVA analysis comparing OSU-03012 treatment to others. Blue: nucleus; green: EdU staining, proliferative marker. Scale bar: 200 µm.

However, the other two PDPK1 inhibitors, GSK2334470 [34, 35] and OSU-03012 [36], failed to demonstrate efficacy. Despite being the most potent and specific PDPK1 inhibitor, GSK2334470 exhibited the least anti-tumor activity in HGSC cell lines (Fig.1D, Fig EV.1B). Meanwhile, OSU-03012, a PDPK1 inhibitor with known secondary target [39], demonstrated cell cytotoxicity in the five-day assay but lacked efficacy in the long-term colony formation assay (Fig.1D, Fig EV.1B). Also, BX-912 quickly induced a dosage-dependent cytotoxic effect 24 hours post-incubation, which was not seen in the OSU-03012 treatment (Fig EV.1C). In a five-day treatment/recovery experiment, BX-912 induced a prolonged cytotoxicity that was not seen in olaparib or OSU-03012 after drug removal (Fig EV.1D). These results suggest that BX-912, despite being a putative PDPK1 inhibitor, may have unique targets contributing to its enhanced efficacy in HGSC cells.

### BX-912 distinctively induces giant multinucleated cell phenotype and cell growth arrest

Along with its anti-tumor cytotoxicity, the BX-912 treatment induced dramatic phenotypic alterations in HGSC cells, characterized by a substantial increase in giant multinucleated cells with flattened cytoskeleton (Fig.1E). Furthermore, the BX-912 treatment led to a reduction in cell growth; this effect was absent from the OSU-03012 treatment, where the surviving cells remained highly proliferative; or from the olaparib treatment, which primarily induced cytotoxicity without suppressing the proliferation of residual cells (Fig.1F). The reduced proliferation was coupled with increased nuclear size following the BX-912 treatment (Fig EV.1E). Furthermore, among the three tested PDPK1 inhibitors, only BX-912 showed significant cytotoxicity across multiple cell lines that have developed resistance to niraparib or olaparib, both are PARP inhibitors [21] (Fig EV.1F). We also found that rapamycin, an inhibitor of AKT/mTOR pathway as the major downstream effector of PDPK1 [40], exhibited minimal effects on HGSC cells (Fig EV.1G). Collectively, these findings suggest that BX-912 employs a unique mechanism of action in killing HGSC cells, likely independent of PDPK1 inhibition.

### PISA assay reveals distinct off-target profiles of PDPK1 inhibitors in HGSC cells

To further investigate the mechanisms driving the phenotypic changes and cytotoxicity induced by BX-912, but not GSK2334470 or OSU-03012, we conducted a PISA assay to identify their potential interacting proteins (Fig.2A). COV362 cells were exposed to drug treatment for a short two-hour period, during which no noticeable changes in cell morphology or viability were observed. The proteomics analysis of the cell lysates identified over 8000 proteins, among which 183, 457, and 339 proteins exhibited significant solubility changes following treatment with GSK2334470, BX-912, and OSU-03012, respectively (P value < 0.05). The PISA results were supported by several previous findings, including DHODH as a secondary target of OSU-03012 [39] and AURKA, a protein linked to cell cycle arrest [41], which was uniquely identified for BX-912 (Fig EV.2A). PDPK1 was detected exclusively in the GSK2334470-treated cells, as GSK2334470 exhibits the highest binding affinity among the tested compounds.

**Figure 2.**
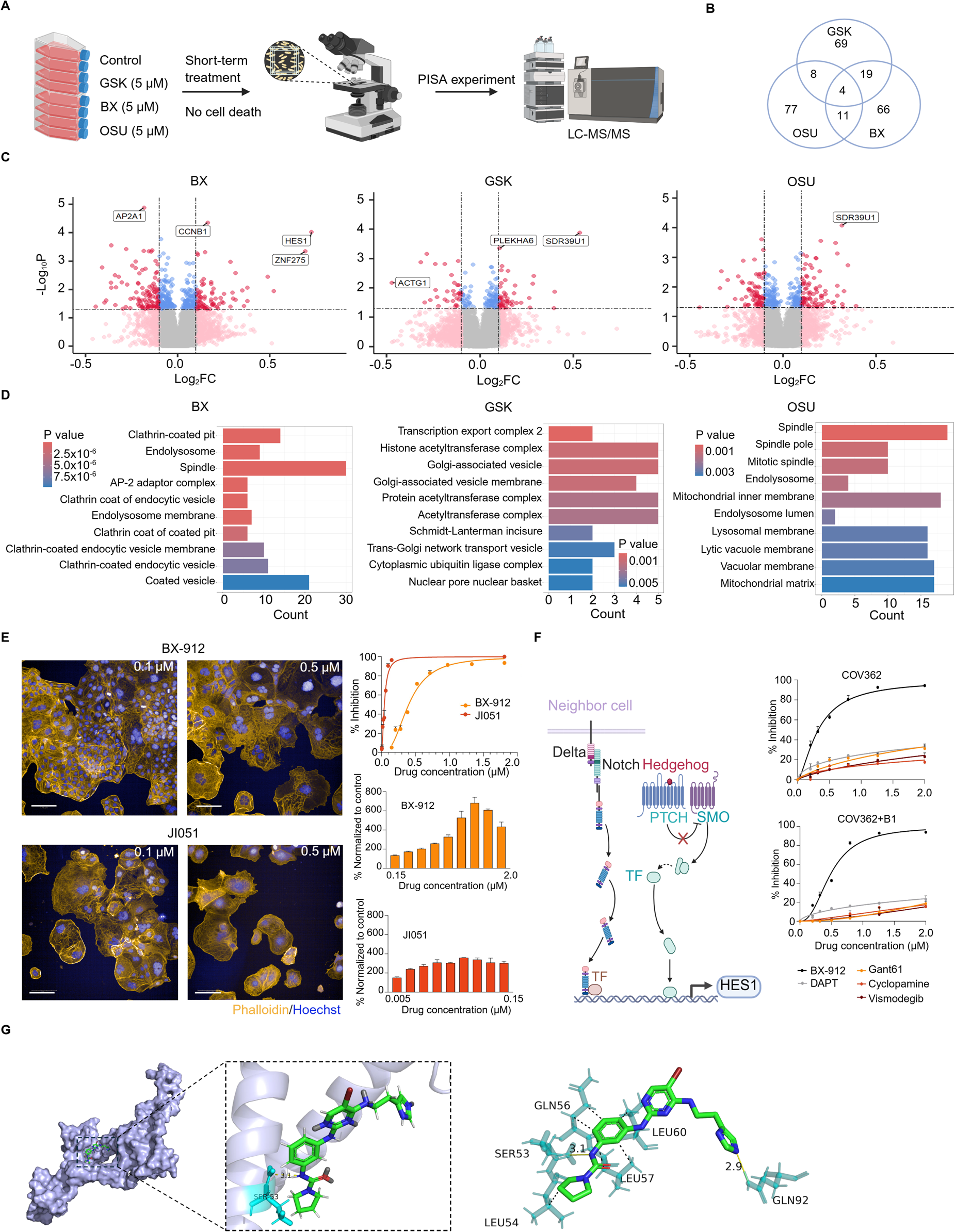
PISA assay reveals the distinct off-target activities of PDPK1 inhibitors. A Workflow of the PISA experiment. B Venn diagram illustrating common and unique proteins identified for each drug (top100, pre-selected by P < 0.05 and ranked by fold change (FC)). C Volcano plots highlighting the most significant protein candidates identified in PISA experiments for each drug, based on P value and FC. Negative and positive FC values indicate respectively decreased and increased thermal stability upon drug interaction, both suggesting structural modifications. D Pathway enrichment (ClusterProfiler) analysis of proteins (P < 0.05) identified in PISA experiments for each drug. E Representative images showing cytotoxicity and morphological changes following BX-912 and JI051 treatment at 0.1 µM and 0.5 µM. Yellow: cytoskeleton, blue: nucleus. Scale bar: 200 µm. Dot plot depicting the drug response curves for BX-912 and JI051, and bar plots showing the nuclear size changes normalized to control. F Schematic representation of classical regulatory mechanisms of HES1 expression. Dose-response curves of multiple inhibitors targeting the Notch (namely DAPT) and Hedgehog (namely GANT61, cyclopamine, vismodegib) pathways are shown in comparison to BX-912. G Docking results for BX-912 and HES1. The zoom in capture shows the top-ranked pose from the 15 ns structure, with BX912 (green sticks) located within the Orange domain pocket. The ligand forms one hydrogen bond with SER53 (yellow dashed line) and engages in multiple hydrophobic and π–π interactions. This pose was found to recur in multiple molecular dynamics-derived conformations, supporting the stability of the binding mode.

Notably, only a limited number of target proteins were shared among the three drugs, suggesting their distinct target interaction profiles (Fig.2B). Transcription factor HES1 emerged as the most significant interacting target for BX-912, not for GSK2334470 or OSU-03012 (Fig. 2C). In contrast, SDR39U1, a member of the epimerase family, ranked as the top interacting protein for both GSK2334470 and OSU-03012. Indeed, molecular docking experiments demonstrated that the binding interactions between SDR39U1 and either OSU-03012 or GSK2334470 were more stable than that with BX-912 (Fig EV.2B).

Subsequent Gene Set Enrichment Analysis (GSEA) analysis identified the PI3K/AKT/mTOR pathway as a common target of all three drugs, supporting the primary mechanism of action of PDPK1 inhibitors (Fig EV.2C). However, when focusing only on statistically significant proteins (P < 0.05), distinct cellular functions were revealed (Fig.2D, Fig EV.2D). These findings suggest that BX-912’s unique efficacy in HGSC may be linked to its off-target activity, particularly involving the transcription factor HES1.

### HES1 is likely the primary target of BX-912 in HGSC cells

HES1 expression exhibits dynamic oscillations during the cell cycle [42]. Indeed, the differential expression of HES1 was observed among individual HGSC cells (Fig EV.2E). Given this heterogeneity, we opted for a pharmacological approach to validate HES1 as a novel target in treating HGSC. We tested JI051, a potent HES1 inhibitor by stabilizing its interaction with the chaperone protein prohibitin 2 (PHB2) outside nucleus [43]. All three HGSC cell lines were sensitive to JI051, comparable to the BX-912 treatment. More importantly, the JI051 treatment also induced the giant multinucleated cell phenotype, similar to BX-912 treatment (Fig.2E).

Previous strategies for HES1 inhibition have been primarily based on targeting its upstream regulatory pathways, such as notch and hedgehog signaling [44–46]. However, in our study, those classic HES1 inhibitors failed to induce cytotoxic effects or phenotypic changes in HGSC cells (Fig.2F). It therefore suggests clarifying alternative regulatory mechanisms that govern the function of HES1 in HGSC is crucial for evaluating its therapeutic potential.

To investigate the interaction between HES1 and BX-912 at the molecular level, we first performed molecular dynamics simulations to equilibrate the HES1 protein, allowing the functional yet flexible WRPW motif to reach a stable conformation (Fig EV.2F). Docking analysis revealed that BX-912 occupies a pocket within the Orange domain of HES1, which is known for dimerization. The top-scoring GOLD pose (ChemPLP score 53.85 and GoldScore 28.06) shows one polar anchoring region and a hydrophobic rim. To characterize the binding mode, Protein-Ligand Interaction Profiler (PLIP) [47] revealed that BX912 formed hydrogen bonds with residues SER53 and GLN92, with donor-acceptor distance of approximately 3.1 and 2.9 Å, respectively. In addition, the ligand forms multiple hydrophobic contacts with Leu54A, Gln56A, Leu57A, and Leu60A (Fig.2G, Fig EV.2G). These hydrogen bonds are expected to stabilize binding within the active pocket, while the hydrophobic interactions create a nonpolar environment that may further support ligand accommodation.

### BX-912 induces a cell cycle arrest in HGSC cells

The multinucleated cells following BX-912 treatment indicate failed cell division; we therefore compared this cellular phenotypic change to the effect of nocodazole, a microtubule-disrupting agent known to disrupt cell cycle progression towards mitosis [48]. Indeed, the multinucleated giant cell phenotype observed after BX-912 and JI051 treatment closely resembled the tetraploid cells generated by nocodazole-induced mitotic arrest (Fig.3A).

**Figure 3.**
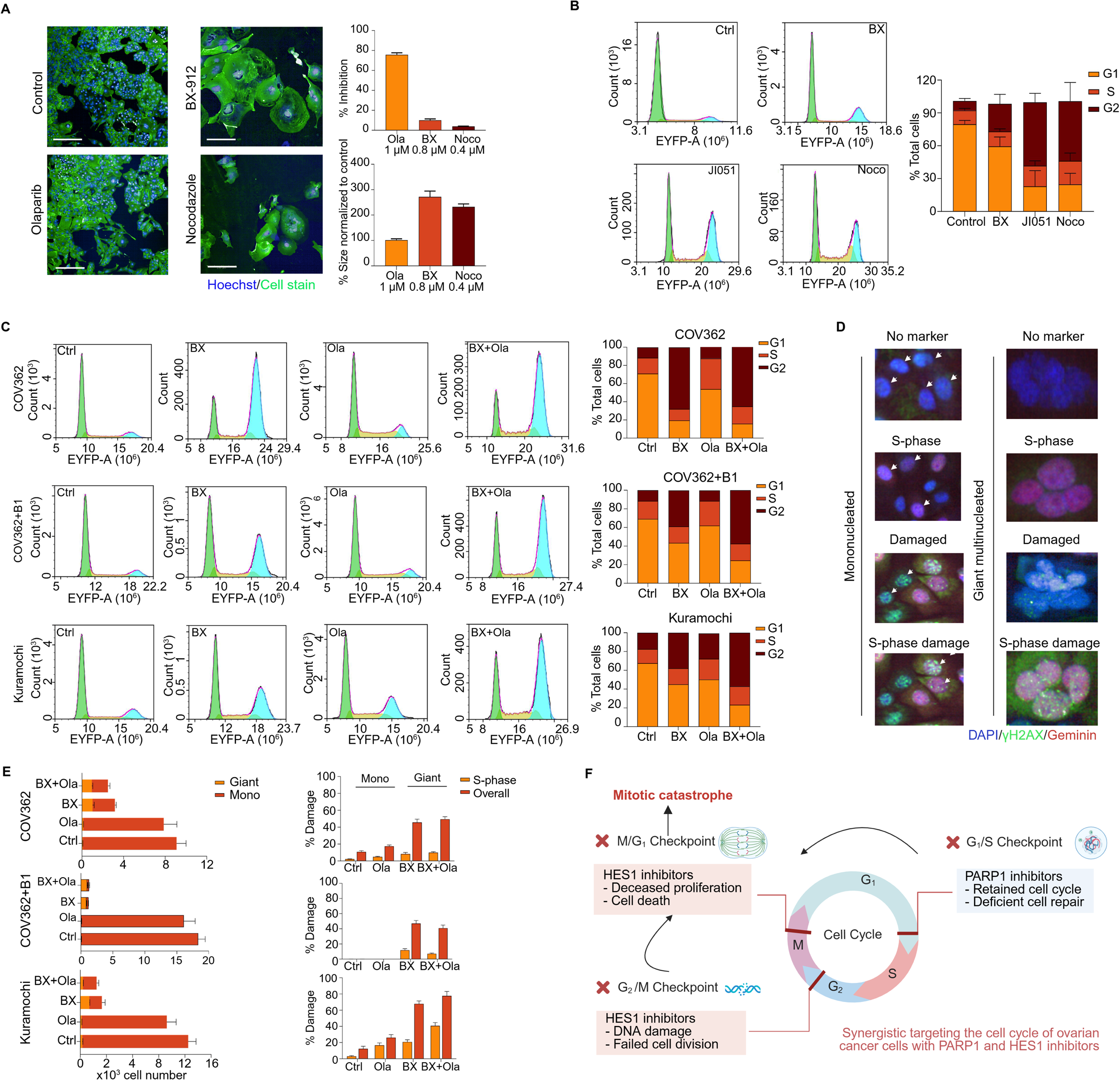
BX-912 induces mitotic catastrophe in HGSC cells. A Representative images and bar plots showing cytotoxicity and phenotypic changes of COV362+B1 cells following the treatment of olaparib (Ola), BX-912 (BX) and the G2-phase blocking agent nocodazole (Noco). Blue: nuclei; green: cell stain. Scale bar: 200 µm. B Cell cycle analysis confirms that the nuclear morphological changes observed after BX-912 treatment result from mitotic arrest. Flow cytometry data are shown for COV362+B1 cells; bar plots represent quantification across three HGSC cell lines following five-day treatment with BX-912, JI051, or nocodazole. C Cell cycle analyses validating the synergy between BX-912 and olaparib in three HGSC cell lines. D Representative immunofluorescent images showing mononucleated cells (indicated by arrows) and giant multinucleated cells. S-phase (S/G2(+M)) cells are geminin-positive (red); damaged cells are with γh2AX foci (green bright dots in the nuclear regions); cells exhibiting S-phase DNA damage are geminin-positive with γh2AX foci. Blue: nuclei. E Bar plots showing the quantification of cell number change. BX-912 treatment leads to accumulation of multinucleated cells with extensive total and S-phase DNA damage. n = 3; data are presented as mean ± SD. F Schematic mode of synergy between olaparib and BX-912: PARP1 inhibitors, such as olaparib, impair homologous recombination repair and subsequently cause S-phase retention. In contrast, BX-912 enhances DNA damage and enforces cell cycle arrest, ultimately driving more HGSC cells—including those that escape PARP1 inhibition—into mitotic catastrophe.

Furthermore, flow cytometry-based cell cycle analysis confirmed that both BX-912 and JI051 induced G2/M phase arrest in HGSC cells, using nocodazole as a reference (Fig.3B). The G2 phase was further prolonged when cells were treated with a combination of olaparib and BX-912 (Fig.3C). On the other hand, olaparib alone increased the proportion of cells in S-phase as expected. Therefore, the addition of BX-912 further impeded cell cycle progression in cells that escape or are resistant to PARP1 inhibition, providing a potential mechanistic explanation for the synergistic effects between BX-912 and olaparib.

### A coordinated damage response and cell cycle arrest leads to mitotic catastrophe by BX-912 together with olaparib

Mitotic catastrophe is a form of cell death resulting from prolonged mitotic arrest and associated DNA damage [49]. Given that mitosis represents a critical phase of the cell cycle susceptible to damage, we assessed the DNA damage status in BX-912-treated HGSC cells by labelling γH2AX and geminin, which is an S/G2 (+M) marker. Using a machine-learning-based image analysis tool [24], we classified post-treatment cells into four distinct cell cycle/DNA damage phenotypes, focusing our quantification on the most distinguishable giant multinucleated and mononucleated categories (excluding binucleated cells) (Fig.3D). Notably, more than 40% of BX-912-induced multinucleated cells exhibited DNA damage, as indicated by γH2AX foci formation, consistent with previous reports linking prolonged mitotic arrest to DNA double-strand breaks [49]. Furthermore, the BX-912 treatment alone led to increased S-phase DNA damage, as evidenced by geminin and γH2AX double positivity—a marker associated with PARP1 inhibitor efficacy [36] (Fig.3E). This increase in S-phase damage was more pronounced in Kuramochi and COV362+B1 cells, which were resistant to olaparib.

These findings suggest that the synergistic cytotoxic effect likely results from BX-912-mediate mitotic arrest and DNA damage induction, coupled with olaparib-induced impairment of DNA repair (Fig.3F).

### BX-912–induced mitotic catastrophe is coordinated by the accumulation of HES1 within the nucleus

Nuclear HES1 abundance is normally maintained through tightly regulated oscillatory expression, a mechanism essential for controlling cell differentiation in both normal and malignant contexts [50, 51]. While the selective HES1 inhibitor JI051 promotes the cytosolic retention of HES1 [43], we observed a pronounced nuclear accumulation of HES1, most evident in multinucleated cells, following BX-912 treatment (Fig.4A, Fig EV.3A). Western blotting further demonstrated that, while both HES1 monomers and dimers were present in the cytosol, only dimers were detected in the nucleus. BX-912 selectively increased nuclear HES1 dimers while depleting cytoplasmic monomers (Fig EV.3B).

**Figure 4.**
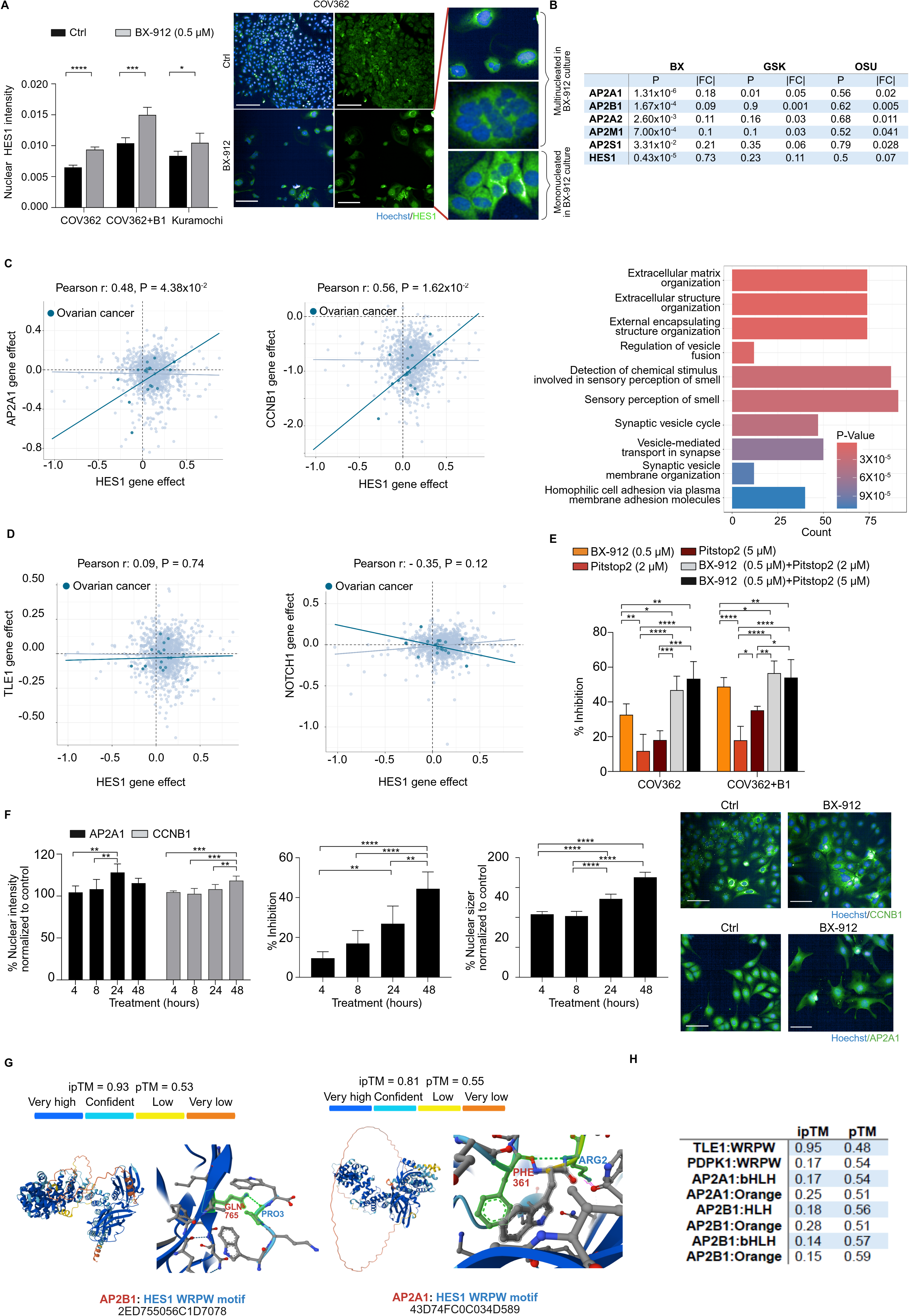
Increased nuclear accumulation of HES1-AP2-CCNB1 protein axis during BX-912-induced mitotic catastrophe. A Left: Mean intensity of HES1 staining measured in nuclear regions after five-day control or BX-912 treatment. **** P < 0.001, *** P < 0.005, * P < 0.05 by t-test. Right: Representative images showing HES1 distribution in control and BX-912-treated cells. Enlarged panels highlight differences in HES1 distribution between mononucleated and multinucleated cells under BX-912 treatment. Blue: nucleus; green: HES1 staining. Scale bar: 200 µm. B Table showing AP2 subunits as BX-912-specific targets in the PISA assay, but not for the other two PDPK1 inhibitors. C Scattered plots showing the co-dependency of AP2A1/HES1 and CCNB1/HES1 for ovarian cancer cell lines; bar plot showing the enriched pathways for genes having significant functional co-dependency with HES1 (P < 0.05 and Pearson correlation > 0.2). D Co-dependency of TLE1/HES1 and NOTCH1/HES1 in ovarian cancer cell lines. E Five-day drug response to BX-912, pitstop2 and their combination in COV362 and COV362+B1 cells. * P < 0.05, ** P < 0.01, *** P < 0.005, **** P < 0.001 by one-way ANOVA test. F Bar plots and representative images showing the increased nuclear intensities of AP2A1 and CCNB1 stainings following a time-course treatment, where paired controls and treated samples were collected at 4-, 8-, 24-and 48-hour following BX-912 treatments. Blue: nucleus; green: CCNB1 or AP2A1 staining. Scale bar: 100 µm. G Structural predictions for binding between AP2B1 and AP2A1 with the WRPW motif of HES1. Predicted binding sites are highlighted in green. The WRPW motif, but not bHLH or Orange domains, shows high interaction potential. Binding between HES1 and TLE1 (via WRPW) is included as a positive control. *pTM* > 0.5 and *ipTM* > 0.8 suggest high-confidence predictions. H Predicted interactions between HES1 domains (bHLH, Orange, WRPW) and AP2 proteins.

Interestingly, our PISA data also highlighted the enrichment of adaptor protein complex 2 (AP-2) subunits and G2-M regulators exclusively in the BX-912-treated condition (Fig.4B, Fig EV.3C). In line with this, clathrin-mediated endocytosis was uniquely targeted by BX-912 (Fig.2D). As HES1 expression showed no correlation with major cell-cycle genes (Fig EV.3C), the short drug–cell exposure time in the PISA assay led us to hypothesize that BX-912 modulates HES1 activity and other top-ranking proteins through a non-transcriptional mechanism.

To explore this unexpected functional connection, we first examined gene co-dependencies using Cancer Dependency Map (DepMap) cell line data. Interestingly, HES1 exhibited stronger co-dependency with AP2A1 and CCNB1 (Fig. 4C) than with TLE1, a well-established transcriptional co-repressor of HES1 [52], or NOTCH1/NUMB, or PHB2, in ovarian cancer cell lines (Fig.4D, Fig EV.3D). Further clusterProfiler analysis of the gene set derived from such DepMap co-dependency data revealed that HES1-associated processes were predominantly linked to vehicle-mediated secretion and transport, despite being a transcription factor (Fig.4C, Fig EV.3E). These findings suggest a novel unrecognized regulatory role of HES1 via the endocytic protein transport machinery.

We further performed a drug perturbation experiment combining BX-912 with pitstop2, an inhibitor known to trap the clathrin-mediated endocytic machinery and is associated with mitotic arrest [53, 54]. While pitstop2 alone had no significant impact on HGSC cell viability, its combination with BX-912 resulted in a further reduction in cell viability compared to BX-912 alone (Fig.4E). Intriguingly, despite being an endocytic protein functioning in the cytosol, AP2A1 was highly enriched in the nucleus, which can be further induced by BX-912 treatment (Fig.4F, Fig EV.3F). Similarly, nuclear CCNB1 levels increased accordingly (Fig.4F), opposing to a transient nuclear transport for initiating the mitotic entry in normal condition [56].

AlphaFold 3 [55] indicated that the WRPW motif, not the bHLH or Orange domains, had strong confidence scores of interaction potential with AP2A1 and AP2B1 (ipTM > 0.6, pTM > 0.5) (Fig.4G), comparable to the known interaction with TLE1 (Fig.4H). It further suggested potential interactions between HES1’s WRPW domain and CCNB1, either directly or mediated through AP2B1 (Fig EV.3G). Overall, these findings propose a model in which BX-912 directly modulates HES1 activity, promoting its dimerization and nuclear translocation together with WRPW-binding partners, thereby contributing to mitotic catastrophe (Fig EV.3H).

### Clinical relevance of targeting HES1 for HGSC treatment

We observed that BX-912 demonstrated potent efficacy across multiple patient-derived HGSC organoids, highlighting the potential of targeting HES1 in this challenging ovarian cancer subtype (Fig.5A). Notably, HES1 expression was significantly elevated in tumor tissues compared to normal ovarian tissues and further increased in metastatic sites (Fig.5B), using the dataset from TNMplot [56]. Additionally, cell-cycle regulator proteins that ranked high in BX-912-PISA list were also more elevated in the malignant ovarian tissues (Fig.5C). In a local HGSC cohort (DECIDER; ClinicalTrials ID: NCT04846933), HES1 expression was consistently detected in tumor epithelial tissues across various sites (Fig EV.4A-B). Its levels were higher in tumor epithelial cells compared to surrounding fibroblasts and immune cells (Fig. 5D). This observation aligns with the greater sensitivity of BX-912 observed in COV362 cells compared to normal epithelial and fibroblast cells (Fig.5E). There was no clear association between HES1 expression and homologous recombination repair status (Fig EV.4C), which was consistent with our observation that BX-912 induced cytotoxicity in all three HGSC cell lines regardless of the HR status. These findings suggest that targeting HES1 is a viable therapeutic strategy in ovarian cancer.

**Figure 5.**
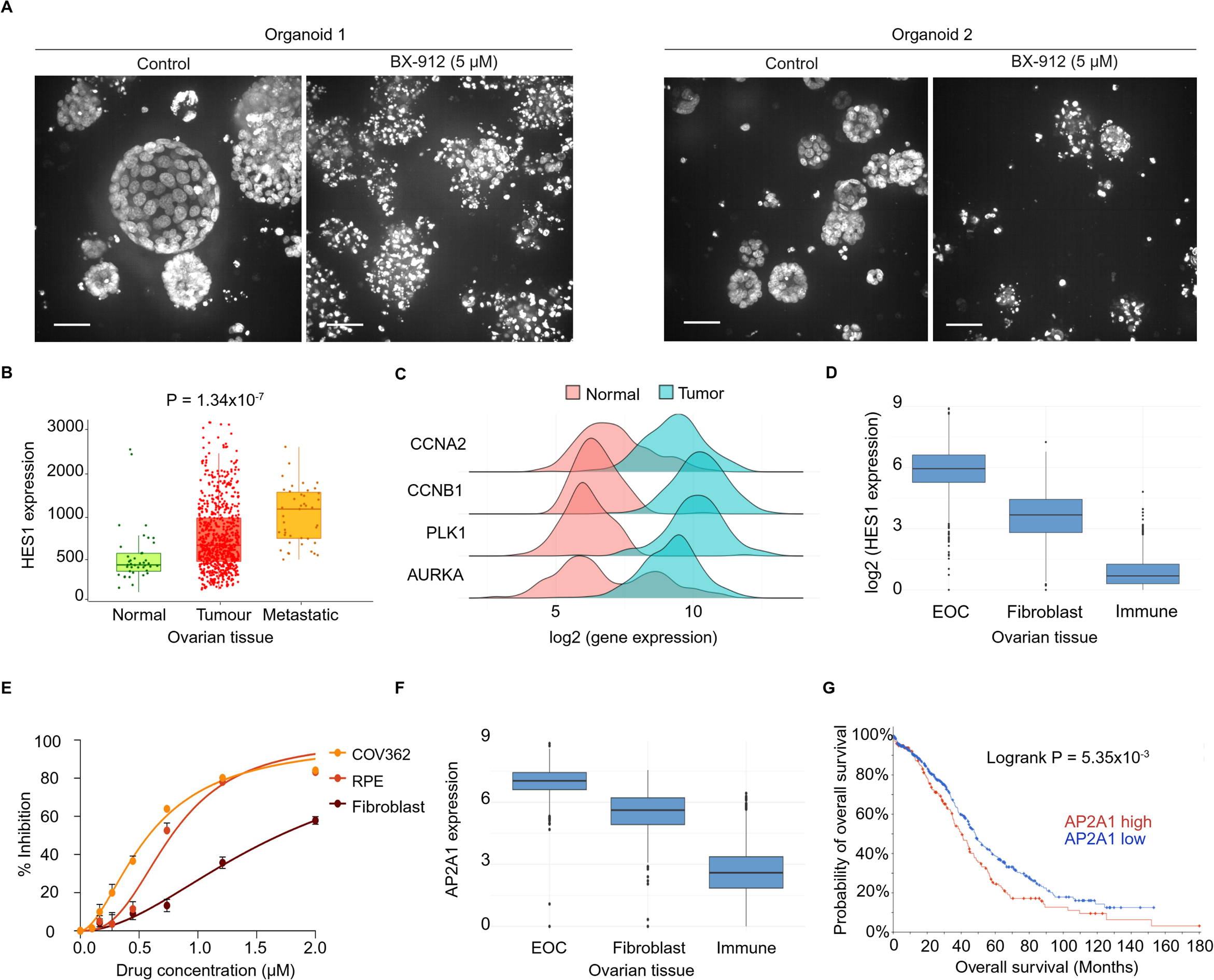
Clinical relevance of targeting HES1 in HGSC. A Representative images showing the efficacy of BX-912 on two HGSC organoids derived from patients. staining: nucleus. Scale bar: 200µm. B Expression of HES1 in normal, tumour and metastatic ovarian tissues. C Expression of G2-M regulators in normal and tumour ovarian tissues. D Expression of HES1 in cancer epithelia (EOC), fibroblast, and immune cells in the DECIDER cohort. E Drug response curves of BX-912 in cancer, normal epithelial (RPE) and 3T3 human fibroblast cells. F Expression of AP2A1 in cancer epithelia (EOC), fibroblast and immune cells in the DECIDER cohort. G Survival analysis of the TCGA-GDC cohort based on AP2A1 expression.

We didn’t detect any clear trend how HES1 expression differed following initial chemotherapy (Fig EV.4D). Furthermore, contrary to a prior study associating low HES1 levels with longer patient survival [57], our analyses of two independent cohorts, TCGA and DECIDER, did not support HES1 expression as a poor prognostic factor for HGSC patient survival (Fig EV.4E).

Derived from the findings of this study, the clinical relevance of adaptor protein AP2A1 was examined within the same HGSC cohorts. AP2A1 was also commonly expressed in tumor epithelial cells from various sites, regardless of HRP/HRD status (Fig EV.4A-C), and its expression was higher in tumor epithelial cells than in immune or fibroblast populations (Fig.5F). While AP2A1 levels showed no consistent changes in post-primary chemotherapy, its expression was negatively associated with patient survival in HGSC cohorts (Fig.5G, Fig EV.4F). Overall, these findings highlight the potential of AP2A1 as a prognostic marker and HES1 as a therapeutic target in ovarian cancer.

## Discussion

In this study, we discovered that the PDPK1 inhibitor BX-912 effectively inhibits the survival and growth of HGSC cells, either alone or together with olaparib, regardless of the HR status. Notably, BX-912 treatment induced a distinctive cellular phenotype characterized by the clustering of multiple nuclei within a single cytoskeleton, an effect absents with other PDPK1 inhibitors and a sign of mitotic catastrophe. Interestingly, the cytotoxic effects of BX-912 were unexpectedly linked to the off-target inhibition of HES1, as identified using the PISA method. PISA and other thermal proteome profiling methods have increasingly been used in various pharmacological studies, primarily for 1) identifying agents with specific cytotoxic effects and their molecular targets [58, 59], 2) profiling the targets of drugs in large-scale screens [60], and 3) validating repurposed drugs by identifying their actual protein targets [61–63]. By identifying a potential protein complex involving CCNB1, HES1, and AP2A1, our study highlights how high-throughput proteomics methods can effectively uncover drug mechanisms of action related to previously underappreciated drug-protein interactions.

Our study demonstrated that BX-912 exerts both anti-proliferative and cytotoxic effects on HGSC cells, with flow cytometry analysis revealing significant G2/M phase arrest. This HES1-induced cell cycle arrest has been considered as irreversible senescence [64], reinforcing HES1 as a promising anti-cancer target. The peak expression of HES1 in the G2 phase suggests its critical role during this stage of the cell cycle [42]. However, intriguingly, we found that inhibiting HES1 expression—either by disrupting the NOTCH1 pathway or using the BET inhibitor, a cytostatic agent that arrests HES1 in the G1 phase [45]—failed to induce cytotoxicity in HGSC cells. This suggests that alternative strategies are needed to effectively silence HES1 function in HGSC.

HES1 has long been proposed as a potential anti-cancer target [51]. Our study, along with findings from [43], demonstrates that effective HES1 inhibition requires detailed characterization of its structural domains and interacting partners. While HES1’s bHLH and orange domains mediate dimerization and regulate subcellular localization [65, 66], AlphaFold 3 predictions reveal that its WRPW motif serves as a critical interface for interactions with both endocytic proteins (AP2A1/AP2B1) and the cell cycle regulator CCNB1. The WRPW domain represents an attractive therapeutic target, as it mediates these newly identified interactions which can compete with binding to the nuclear co-repressor TLE1; however, its intrinsic disordered nature may pose challenges for direct drug targeting. In contrast, the Orange domain, which likely harbors the BX-912 binding pocket, remains stably folded under all conditions [67]. Elucidating how HES1 dimerization influences protein binding and nuclear translocation could provide a framework for developing novel strategies to target transcription factors like HES1 that lack conventional druggable pockets [68].

We also observed that BX-912 directly induces DNA damage in HGSC cells. HES1 is known to play a role in regulating DNA damage and repair processes. In B-cell acute lymphoblastic leukemia, an interaction between PARP1 and HES1 inhibits HES1 function, leading to PARP1 activation and apoptosis, partially via the notch pathway [69]. Additionally, BRCA1 has been shown to interact with HES1 to support interstrand crosslink DNA repair, complementing PARP1-mediated homologous recombination repair [70]. HES1 has also been identified as a binding partner of REV7 in the Fanconi anemia core complex [71]. In our study, BX-912-induced multinucleated cells exhibited extensive DNA damage that was not restricted to the typical S-phase damage caused by PARP inhibitors. DNA damage response and cell cycle arrest are closely linked [72], and our findings suggest that BX-912 treatment can trigger both events, ultimately leading to cell death. Further investigation into the orchestration of DNA damage and cell cycle arrest in mitotic catastrophe by HES1 may provide valuable insights into the interconnection of those cancer hallmark events [73].

The clinical features and therapeutic responsiveness of cancer are primarily driven by dysfunctional transcriptional programs in cancer cells [74]. However, transcription factors have been challenging to target for therapeutic purposes [75]. HES1, a transcription factor implicated in cancer progression, shows promise as a potential therapeutic target based on our study and previous findings [76, 77]. A primary concern in targeting transcription factors is the risk of general cytotoxicity and severe side effects. Notably, our study showed that HES1 expression is the highest in cancer cells in comparison to immune and stromal compartments. Additionally, previous research in intestinal tissue demonstrated that HES1 exerts distinct regulatory effects in normal versus tumor cells, suggesting its potential as a tumor-specific therapeutic target [78]. The observed dose-dependent induction of mitotic catastrophe by BX-912 in our study highlights the possibility of using lower doses to reduce treatment toxicity, selectively inducing mitotic catastrophe at levels tolerated by normal epithelial and other cell types within the tumor environment.

## Methods

### Cell culture

COV362 cells (HR-deficient, Merck, 07071910) and derived COV362+BRCA1 cells (COV362+B1 in this manuscript, where the HR-deficiency was reversed by re-expressing *BRCA1*) [20] were cultured in Dulbecco’s Modified Eagle Medium (DMEM, high glucose, pyruvate, no glutamine, Gibco 21969035) supplied with 10% fetal bovine serum (FBS) (Gibco, 10270106), 1x penicillin–streptomycin (P.S.) (Gibco, 15140122), and 1x GlutaMAX (Gibco, 35050038). Kuramochi cells (JCRB Cell Bank, JCRB0098) were grown in RPMI1640 medium (Thermo Fisher, 31870025) supplied with 10% FBS, 1x GlutaMAX, and 1x P.S. All cell lines were authenticated.

Niraparib-resistant (NA1 and NA3) and olaparib-resistant (OA5) cells were developed from *TP53* and *BRCA1*-deficient cells from RPE-1 hTERT cells [21]. These cells were cultured in DMEM and supplied with 10% FBS, 1x P.S., and 1x GlutaMAX. The cross-resistance to olaparib was confirmed before testing with BX-912.

HGSC organoid samples were cultured in Cultrex RGF BME type 2 (Bio-techne, 3533-010-02) as previously described [22]. Before adding BX-912, a 10-day culture time allowed the formation of organoids.

### Small molecule inhibitors

Olaparib (HY-10162), BX-912 (HY-11005), OSU-03012 (HY-10547), GSK2334470 (HY-14981), JI051 (HY-117113), Nocodazole (HY-13520), rapamycin (Sirolimus, HY-10219), DAPT (HY-13027), GANT61 (HY-13901), Cyclopamine (HY-17024), Vismodegib (HY-10440) and pitstop 2 (HY-115604), were all purchased from MedChemExpress.

### Drug combination screening and synergy scoring

For the high-content drug combination assays, Kuramochi cells were seeded at a density of 2,500 cells per well in 96-well black clear-bottom plates (Corning, 3631) or 1,000 cells per well in 384-well black clear-bottom plates (Corning, 3764). COV362 and COV362+B1 cells were seeded at 1,500 cells per well in 96-well plates and 800 cells per well in 384-well plates, respectively. After overnight incubation to allow cell attachment, small molecule inhibitors were administered using the automated D300e Digital Dispenser (TECAN). Following a five-day treatment period, cell viability was assessed through image-based analysis. Drug combination synergy scores were calculated using SynergyFinder Plus [23].

### Immunofluorescent staining and image analysis

At the endpoint of drug treatment, cells were fixed with freshly prepared 4% paraformaldehyde (Sigma-Aldrich, 158127) in phosphate-buffered saline (PBS) for 30 minutes, followed by three washes with PBS containing 0.1% bovine serum albumin (BSA; Sigma-Aldrich, A7906). For antibody staining requiring permeabilization, cells were treated with 0.1% Triton X-100 (Merck Millipore, 9002-93-1) for 25 minutes and subsequently blocked in PBS containing 3% BSA for 30 minutes.

The fixed and permeabilized cells were incubated overnight at 4°C with 1 :1000 dilution of anti-HES1 rabbit polyclonal antibody (Proteintech, PA5-28802), 1:1000 dilution of anti-CCNB1 (Proteintech, 29887-1-AP and 1:1500 dilution of anti-AP2A1 rabbit polycolonal antibody (Proteintech, 55004-1-AP in PBS with 3% BSA. The following day, cells were washed and incubated with Alexa Fluor™ 488 goat anti-rabbit secondary antibody (Thermo Fisher Scientific). Cytoskeleton visualization was achieved using Alexa Fluor™ 555 Phalloidin (Thermo Fisher Scientific) for 30 minutes. Nuclei were counterstained with 1 µM Hoechst (Thermo Fisher Scientific).

Whole-well imaging was performed using the Opera Phenix high-content microscope (PerkinElmer). Quantification of cell numbers and extraction of phenotypic information was conducted with an analysis pipeline developed in CellProfiler (v4.2.6), employing object segmentation and area extraction methods.

### DNA damage analysis

Cultured cells were fixed in 4% paraformaldehyde and stored overnight at 4°C before immunofluorescence (IF) staining. The cells were first permeabilized with PBS/0.2% Triton X-100, followed by washes and blocking with normal donkey serum (NDS). They were then incubated overnight at 4°C with primary antibodies, including mouse monoclonal anti-γH2A.X (phospho S139) (Abcam, ab22551; 1:1000 dilution) and rabbit monoclonal anti-geminin (Abcam, ab195047; 1:500 dilution). The following day, the cells were washed and incubated with corresponding secondary antibodies, Alexa Fluor™ 488 Donkey anti-mouse and Alexa Fluor™ 647 Donkey anti-rabbit (both from Thermo Fisher Scientific), followed by nuclear counterstaining with Hoechst. Finally, the stained cells were visualized using the Opera Phenix high-content microscope (PerkinElmer). To analyze phenotype-specific cell subpopulations, the software Advanced Cell Classifier (ACC) was employed following the user manual [24].

### Cell cycle analysis

After five days of drug treatment, HGSC cells were harvested and washed twice with ice-cold PBS before being dropwise fixed in ice-cold 70% ethanol. The fixed cells were stored at 4°C for at least four hours. To analyze cell cycle phases, FxCycle™ PI/RNase Staining Solution (Thermo Fisher Scientific) was used following the manufacturer’s protocol. Briefly, cell pellets (∼1 × 10⁶ cells per sample) were resuspended in 0.5 mL FxCycle™ PI/RNase Staining Solution. Samples were incubated in the dark for 30 minutes at room temperature and analyzed using flow cytometry with 488-nm excitation and emission collected using a 585/42 bandpass filter.

### Colony formation assay

HGSC cell lines were seeded in 6-well plates at a density of 500 cells per well. After overnight incubation to allow cell attachment, drugs or the corresponding vehicle control (DMSO) were manually added. The culture medium was refreshed every five days with fresh media containing the respective drugs until the three-week endpoint.

For recovery experiments, after one week of drug treatment, cells were washed twice with normal culture media and cultured in drug-free media for an additional week. To fix the colonies, cells were washed twice with ice-cold PBS and fixed in ice-cold 100% methanol for 15 minutes on ice. After fixation, methanol was aspirated, and 2 mL of 0.5% crystal violet solution (500 mg Crystal Violet, 25 mL methanol, 75 mL water) was added to each well for 10 minutes. Excess crystal violet was removed, and the wells were washed until no residual dye remained. Plates were dried overnight and stored at room temperature.

### Western Blot

After the drug treatments, cells were harvested from 10 cm petri dishes using 0.05% trypsin (diluted from 0.25% Trypsin-EDTA, Gibco). The cell pellets were washed two times with ice-cold PBS with 1x Halt protease inhibitor (Thermo Fisher Scientific). To collect the cytosolic proteins, the pellets were suspended and incubated in ice cold nuclear extraction buffer (10 mM Tris-HCL, pH 7.4, 10 mM NaCl, 3 mM MgCl2, 0.5% (v/v) NP-40 (IGEPAL) for 10 min before centrifuging at 1500 rpm for 5 min at 4°C. Cytosolic proteins were therefore collected in the supernatant. To obtain the nuclear proteins, the nuclear pellets were re-suspended in 1x hot lysis buffer (consisting of 62.5 mM Tris, pH6.8; 2% SDS, 10% Glycerol and 0.002% Bromophenol blue). After a short sonification, cell lysates were obtained following 10 min of centrifugation at 10,000x g. After running SDS-page gels, western blotting was performed using trans-blot turbo transfer system (Bio-Rad). The obtained blot was blocked for one hour in 2% BSA with 0.1% tween and incubated with anti-HES1/anti-AP2A1/anti-CCNB1 antibody (1:1000) overnight. The blots were washed and incubated in IRDye® 800CW Goat anti-Rabbit IgG (H + L) (1:10000, LICORbio^TM^) for 30 min. To measure the protein-loading, anti-alpha tubulin antibody [DM1A] (abcam, ab7291, diluted 1:1000) was used. Images were obtained with Odyssey® imaging system (LICORbio^TM^) and the band intensity was measured with ImageJ/FIJI.

### PISA analysis

Cell pellets were resuspended in ice-cold PBS containing protease and phosphatase inhibitor cocktails (Roche). Thermal treatment was conducted following a published protocol with minor modifications [9]. Briefly, cells were aliquoted into a PCR plate and subjected to a 3-min thermal treatment across ten different temperatures: 43.9°C, 44.9°C, 46.5°C, 48.5°C, 50.5°C, 52.5°C, 54.5°C, 56.5°C, 58.1°C, and 59.1°C. Subsequently, plates were incubated for 6 minutes at room temperature. Cell lysis was achieved through five cycles of freezing in liquid nitrogen and thawing at 35°C. Equal aliquots from each temperature point were combined, and Nonidet P-40 substitute was added to a final concentration of 0.4%. Samples were incubated at 4°C for 1 hour with gentle agitation at 350 rpm, followed by ultracentrifugation at 149,000 × g for 30 minutes at 4°C. Supernatants were collected, and proteins were reduced with 20 mM DTT and alkylated with 40 mM iodoacetamide. Proteins were precipitated overnight with ice-cold acetone (1:6 v/v). Pellets were dissolved in 8 M urea in 50 mM TEAB, with the urea concentration subsequently reduced to 2.7 M before digestion with Lys-C (Fujifilm Wako) for 6 hours at 30°C with shaking at 350 rpm. Urea concentration was further diluted to 0.8 M by adding 50 mM TEAB, and proteins were digested overnight with sequencing-grade modified trypsin (Promega). Digested peptides were desalted using Sep-Pak tC18 100 mg 96-well plates. Peptide concentrations were measured using a Qubit Flex fluorometer (Invitrogen), and 50 µg of each sample was labeled with the TMTpro 16plex Label Reagent Set (Thermo Scientific).

TMTpro-labeled samples were fractionated offline using an Agilent 1260 Infinity II HPLC system equipped with an XBridge Peptide BEH C18 column (300 Å, 3.5 µm, 2.1 mm × 250 mm; Waters) at a flow rate of 200 µL/min and maintained at 30°C. The gradient elution program followed [10], and the collected fractions were pooled into 16 samples.

Liquid chromatography-electrospray ionization tandem mass spectrometry (LC-ESI-MS/MS) analyses were performed on an Easy-nLC1200 system (Thermo Scientific) coupled to an Orbitrap Exploris 480 mass spectrometer (Thermo Scientific) equipped with a nano-electrospray ionization source and FAIMS Pro Duo interface (Thermo Scientific). Compensation voltages of -40 V, -60 V, and -80 V were applied. Peptides were first loaded onto a trapping column and then separated on a 15 cm C18 analytical column (75 µm × 15 cm, ReproSil-Pur 3 µm 120 Å C18-AQ; Dr. Maisch HPLC GmbH). The mobile phases consisted of 0.1% formic acid in water (solvent A) and 80% acetonitrile with 0.1% formic acid (solvent B). A 120-minute gradient was employed: 7% to 24% solvent B over 62 minutes, 24% to 39% solvent B over 48 minutes, followed by an increase to 100% solvent B over 5 minutes, and a 5-minute wash with 100% solvent B.

Mass spectrometry data were acquired using Thermo Xcalibur 4.6 software (Thermo Fisher Scientific). A data-dependent acquisition (DDA) method was utilized, consisting of an Orbitrap MS survey scan over a mass range of 350–1750 m/z with a resolution of 60,000 and a maximum injection time of 50 ms. This was followed by higher-energy collisional dissociation (HCD) fragmentation with a resolution of 45,000 and a maximum injection time of 120 ms for the most intense peptide ions, operating in top-speed mode with a cycle time of 1 second for each compensation voltage.

Data files were processed using Proteome Discoverer 3.0 software (Thermo Scientific) with the CHIMERYS search engine. Searches were conducted against the SwissProt database (v2023_1) filtered for *Homo sapiens* entries. Carbamidomethylation of cysteine, TMTpro modification of lysine, and TMTpro modification of peptide N-termini were set as static modifications, while methionine oxidation was specified as a variable modification. Quantitation at the protein level was based solely on unique peptides. Abundance values for peptides and proteins were determined from the intensities of TMTpro reporter ions.

### Molecular docking experiment

Three-dimensional structures of proteins and ligands were obtained from the RCSB Protein Data Bank and PubChem, respectively. RCSB: PDPK1-2XCH; SDR39U1-4B4O. To ensure sidechain accuracy, only protein structures determined by X-ray crystallography with a resolution better than 2 Å were selected. Following preprocessing steps, such as adding hydrogen atoms, docking simulations were performed using AutoDock Vina. During the docking process, a cubic grid box with dimensions of 20 Å per side was employed, and all other parameters were set to their default values.

For specifically docking BX-912 to HES1, we first obtained the initial HES1 protein structure from AlphaFold Protein Structure Database (UniProt ID: Q14469) and removed irrational conformations by Molecular Dynamics (MD) simulations of 35ns at 300K using Gromacs 2025.2 [25]. The binding site was defined around the disordered but functionally-annotated WRPW motif (Trp276–Arg277–Pro278–Trp279). After an initial structural adjustment within the first 8ns, the system reaches a stable plateau at approximately 2.0 nm. The RMSD remains relatively constant with only minor fluctuations, reflecting natural backbone flexibility rather than structural drift. This stability confirms that the protein conformation is well-equilibrated, making it suitable for use as a representative structure in subsequent molecular docking studies. Five receptor conformations from MD snapshots (15, 20, 25, 30, 35 ns) were used for ensemble docking with CCDC GOLD 2024.2.0 [26].

### Clinical data analysis

The DECIDER cohort is a prospective, longitudinal study involving HGSC patients treated with standard-of-care therapy at Turku University Hospital (ClinicalTrials.gov identifier: NCT04846933). Bulk RNA sequencing (RNA-seq) reads from this cohort were processed using the SePIA pipeline [27] within Anduril 2 [28], as previously described [29, 30]. Low-quality bases were trimmed with Trimmomatic [31] v0.33, and the resulting reads were aligned to the GRCh38.d1.vd1 reference genome with GENCODE v25 annotations using the STAR aligner [32] v2.5.2b, allowing up to 10 mismatches. Gene-level effective counts were quantified with eXpress [33] v1.5.1, and the batch-effect correction was performed using the POIBM [34]. Bulk RNA-seq data decomposition was conducted with the PRISM framework [35], enabling the extraction of sample composition, scale factors, and cell-type-specific whole-transcriptome profiles. The RNA expression matrix was normalized by dividing expression values by adjusted cell-type proportions, which were scaled using gain values, log-transformed, and summarized as sample-wise means.

The full cohort included 1159 samples. For the survival analysis, only treatment-naive, solid, representative tissue samples were included. One sample per patient was selected based on the highest tumor fraction, resulting in a subset of 262 samples, for which 251 samples have survival information. The normalized cancer-specific component derived from PRISM decomposition was used for the analysis. To determine an optimal threshold for stratifying patients into high-and low-expression groups, the maxstat R-package (v0.7-25) was employed to perform a maximum log-rank statistic test. Kaplan-Meier survival curves were then generated using the survminer R-package (v0.5.0).

For tissue-specific gene expression analyses, sample selection criteria similar to those used in the survival analysis were applied. The primary HGSC sample collection sites included the adnexa (adn), mesentery (mes), omentum (ome), ovary (ova), peritoneum (per), and fallopian tube (tub). Sites with fewer than five samples were excluded, resulting in a total of 260 samples. The analysis of HR status was done following the previously published protocols [36, 37], samples without HR-status information and tissue types with fewer than five samples in either the HR-proficient (HRP) or HR-deficient (HRD) groups were excluded, resulting in 229 samples. Only sample pairs from the same tissue site were included when analyzing expression changes across treatment phases. Plots were generated for omental samples, where 37 matching pairs were identified.

### Statistical analysis

All data are presented with at least three biological replicates per experiment (mean, unless otherwise specified in the figure legends. Statistical significance is indicated in each legend, with p < 0.05 denoting a statistically significant difference. Genes were annotated using the org.Hs.eg.db (v3.15.0) package and enriched and analyzed using the clusterProfiler (v4.4.4) package, GSEA (v4.3.3). Other plots and analyses were generated using GraphPad Prism 10.

## Abbreviations

HGSC: Ovarian high-grade serous cancer
PDPK1: phosphoinositide-dependent kinase 1
PISA: Proteome Integral Solubility Alteration
CCNB1: G2/mitotic-specific cyclin-B1
TPP: thermal protein profiling
HRP: HR-proficient
HRD: HR-deficient

## Acknowledgements

Proteomics analyses were performed at the Turku Proteomics Facility, University of Turku and Åbo Akademi University. The facility is supported by Biocenter Finland.

FIMM High Content Imaging unit services were used for imaging analysis.

## Author contribution

JB and JT designed the project; JB conducted the project and wrote the manuscript; SP, AA and EK performed DNA damage experiment supervised by LK; JD performed flow cytometry experiments; JH provided the samples and clinical parameters for analyses.; SL performed clinical data analysis supervised by SH; WY performed pathway enrichment analyses; MP and OK (Turku Proteomic Core Facility) performed PISA experiment; DM performed drug response study together with JB; CL performed protein docking (SDR39U1 and BX/OSU/GSK) experiment supervised by ML; WH performed docking between BX-912 and HES1. JE, JH, AV, SH, LK and JT reviewed the manuscript.

---

**Figure EV1.**
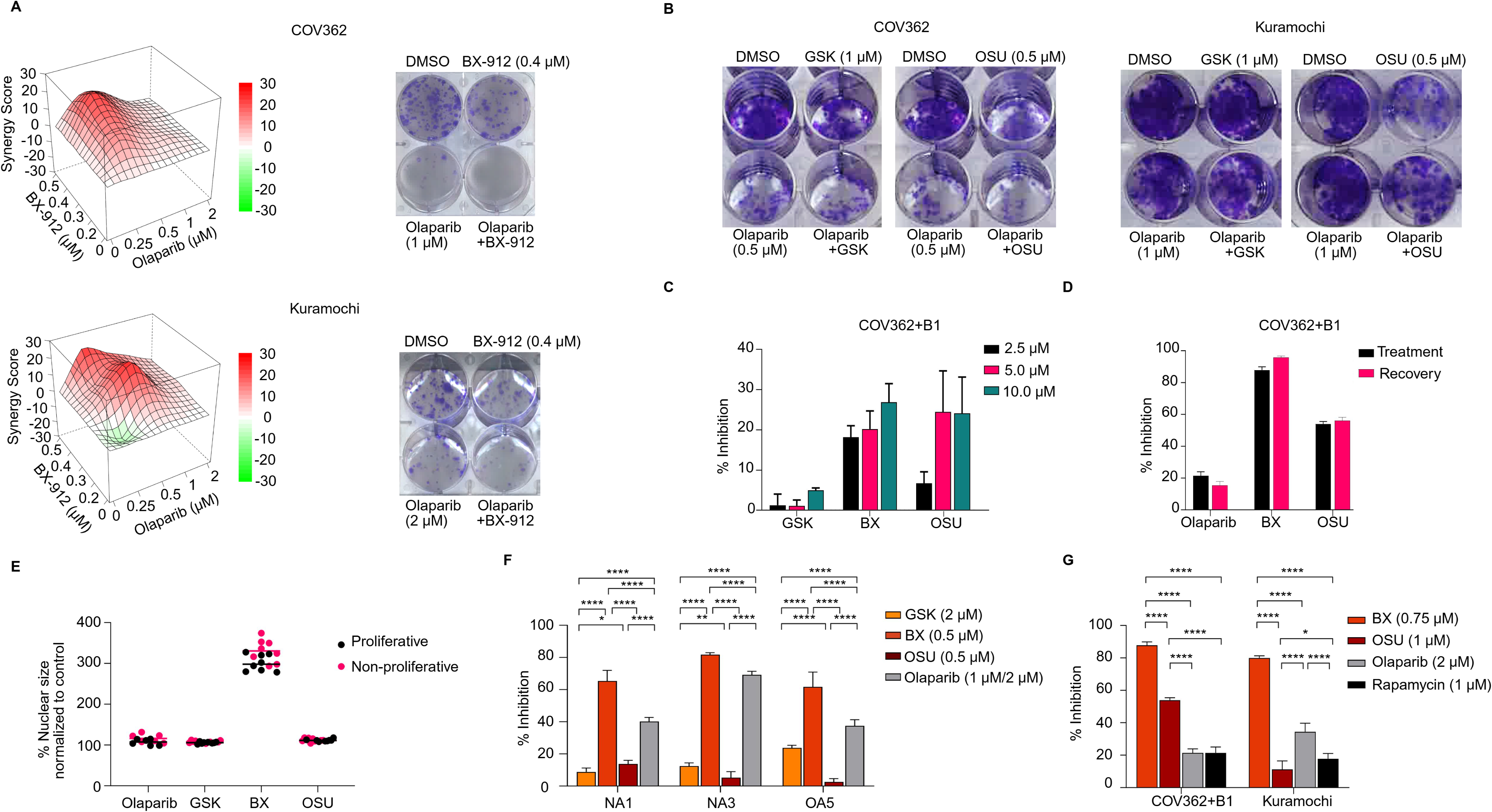
Off-target effects of BX-912 contribute to its cytotoxicity. A Drug synergy matrices and long-term colony formation assays demonstrate the synergistic effects of BX-912 and olaparib in COV362 and Kuramochi cell lines. B Long-term colony formation assays showing the outcomes of olaparib + OSU-03012 and olaparib + GSK2334470 treatments. C Cytotoxic effects of three PDPK1 inhibitors in a concentration-dependent manner at 24 hours post-treatment in COV362+B1 cells. D Cytotoxic effects of olaparib (1 µM), BX-912 (1 µM), and OSU-03012 (1 µM) five days after drug removal, following an initial five-day treatment. E Relationship between nuclear size and proliferation rate after drug treatment. F Efficacy of three PDPK1 inhibitors across two niraparib-resistant and one olaparib-resistant cell lines. * P value < 0.05, ** P value < 0.01, **** P value < 0.001 by one-way ANOVA. G Effects of five-day treatments with PDPK1 inhibitors, olaparib, and the mTOR inhibitor rapamycin. * P value < 0.05, ** P value < 0.01, **** P value < 0.001 by one-way ANOVA.

**Figure EV2.**
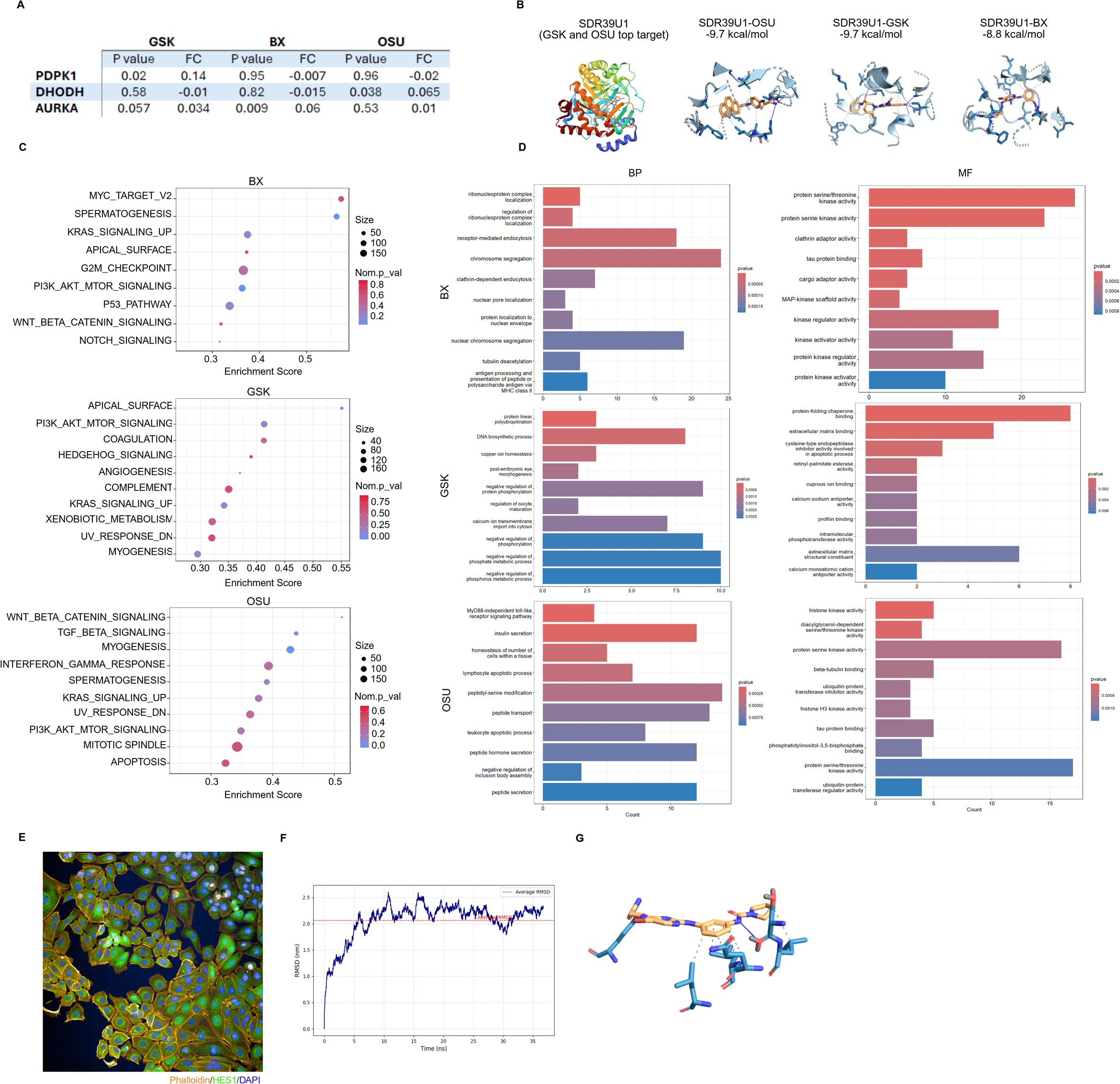
HES1 as the primary target of PDPK1 inhibitor BX-912. A Known off-targets and their PISA results for the three PDPK1 inhibitors. B Molecular docking highlights the interactions between SDR39U1 (ranked as the top target for GSK2334470 and OSU-03012) with the three PDPK1 inhibitors. Lower binding energy values indicate more stable interactions. C-D Pathways enriched by all identified proteins (C) and statistically significant proteins (P < 0.05) (D) in the PISA assays. E Representative images showing heterogeneous expression of HES1 in HGSC cells. Blue: nucleus; Yellow: cytoskeleton; Green: HES1. F HES1 molecular dynamics simulations to acquire representative structure for subsequent docking experiments. After an initial structural adjustment within the first 8ns, the system reaches a stable plateau at approximately 2.0 nm, with only minor fluctuations, reflecting natural backbone flexibility rather than structural drift. This stability confirms that the protein conformation is well-equilibrated, making it suitable for use as a representative structure in subsequent molecular docking studies. Five receptor conformations from MD snapshots (15, 20, 25, 30, 35 ns) were used for ensemble docking. G Bond highlighting: The top-scoring GOLD pose shows one polar anchoring region and a hydrophobic rim. PLIP (4) detected two hydrogen bonds: (i) the ligand donates to Ser53A O2 (H–A 2.16 Å; D–A 3.11 Å; angle 156.6°), and (ii) Gln92A donates to the ligand N2 (H–A 1.87 Å; D–A 2.88 Å; angle 176.3°). In addition, the ligand forms multiple hydrophobic contacts with Leu54A, Gln56A, Leu57A, and Leu60A (closest heavy-atom distances ∼3.3–4.0 Å). Together, these interactions are consistent with a well-packed pose stabilized by specific H-bonds and a leucine-rich hydrophobic environment.

**Figure EV3.**
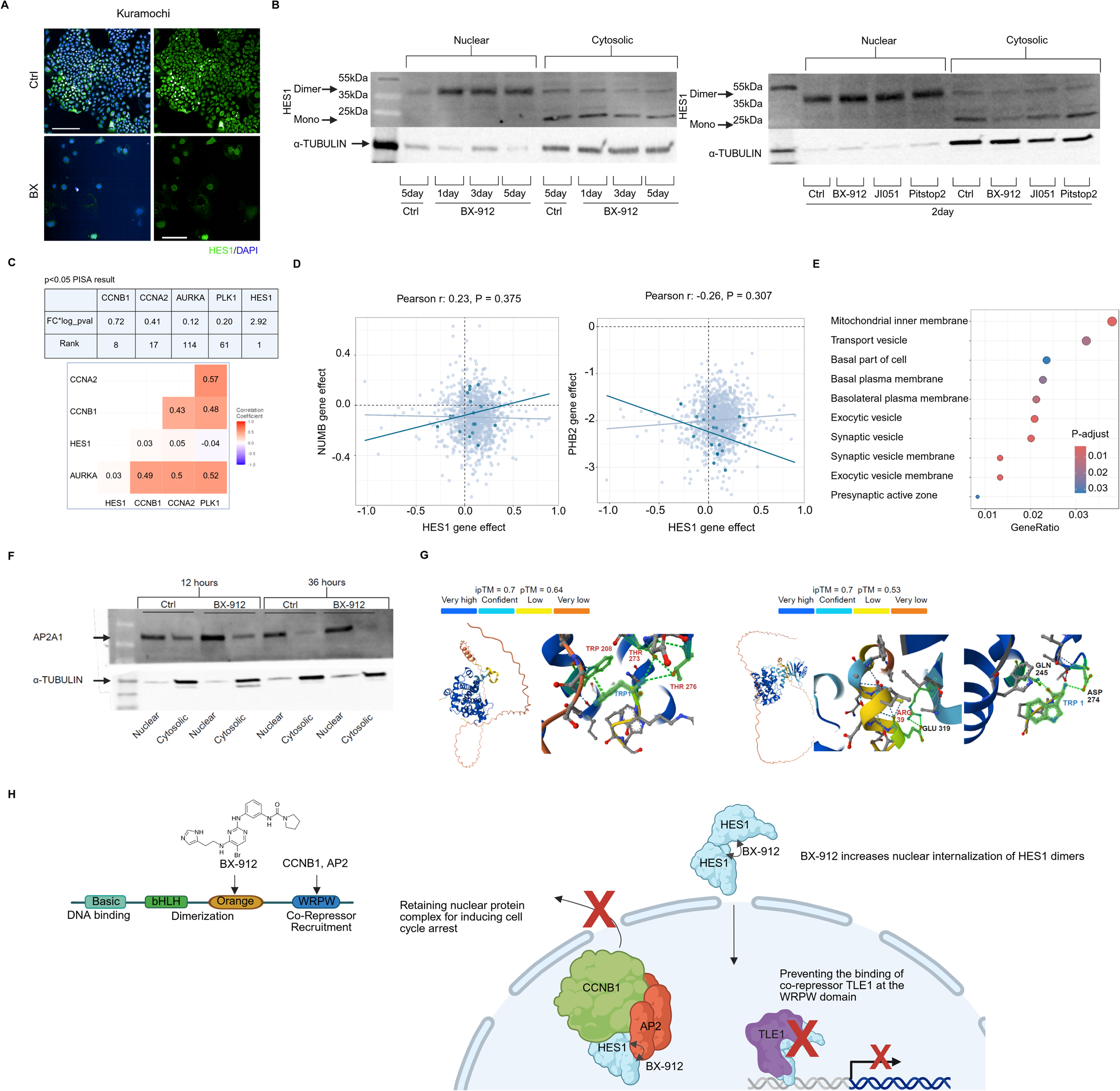
Increased nuclear accumulation of multiple BX-912-PISA top proteins following BX-912-induced mitotic catastrophe. A Representative images showing HES1 distribution in control and BX-912 treated KURAMOCHI cells. Blue: nucleus, green: HES1 staining. Scale bar: 200 µm. B Western blot images showing the time-course changes in HES1 nuclear and cytosolic protein levels following BX-912 treatment (left), and after 48-hour treatment with BX-912, JI051, pitstop2 or control (right). Two bands were detected in cytosolic samples, the lower of which may represent monomeric HES1. C Top: PISA result rankings of multiple cell cycle regulators. Bottom: co-expression correlations between G2 cell cycle regulators and HES1 in ovarian tumor tissues (TCGA dataset). D Co-dependency between HES1 and its known regulators NUMB and PHB2 (Depmap). E GSEA analysis of proteins showing significant co-dependency with HES1 (P < 0.05). F Western blot images showing the time-course changes in AP2A1 nuclear and cytosolic protein levels following 12-hour and 36-hour BX-912 treatment, paired controls were included. G AlphaFold 3 predictions of structural interactions between CCNB1 and AP2B1-CCNB1 with the WRPW motif of HES1. H Proposed model of BX-912–induced mitotic catastrophe. BX-912 binds the Orange domain of HES1, enhancing dimerization and nuclear localization, while the WRPW motif engages CCNB1 and AP-2 proteins and competes with TLE1, collectively disrupting cell cycle progression.

**Figure EV4.**
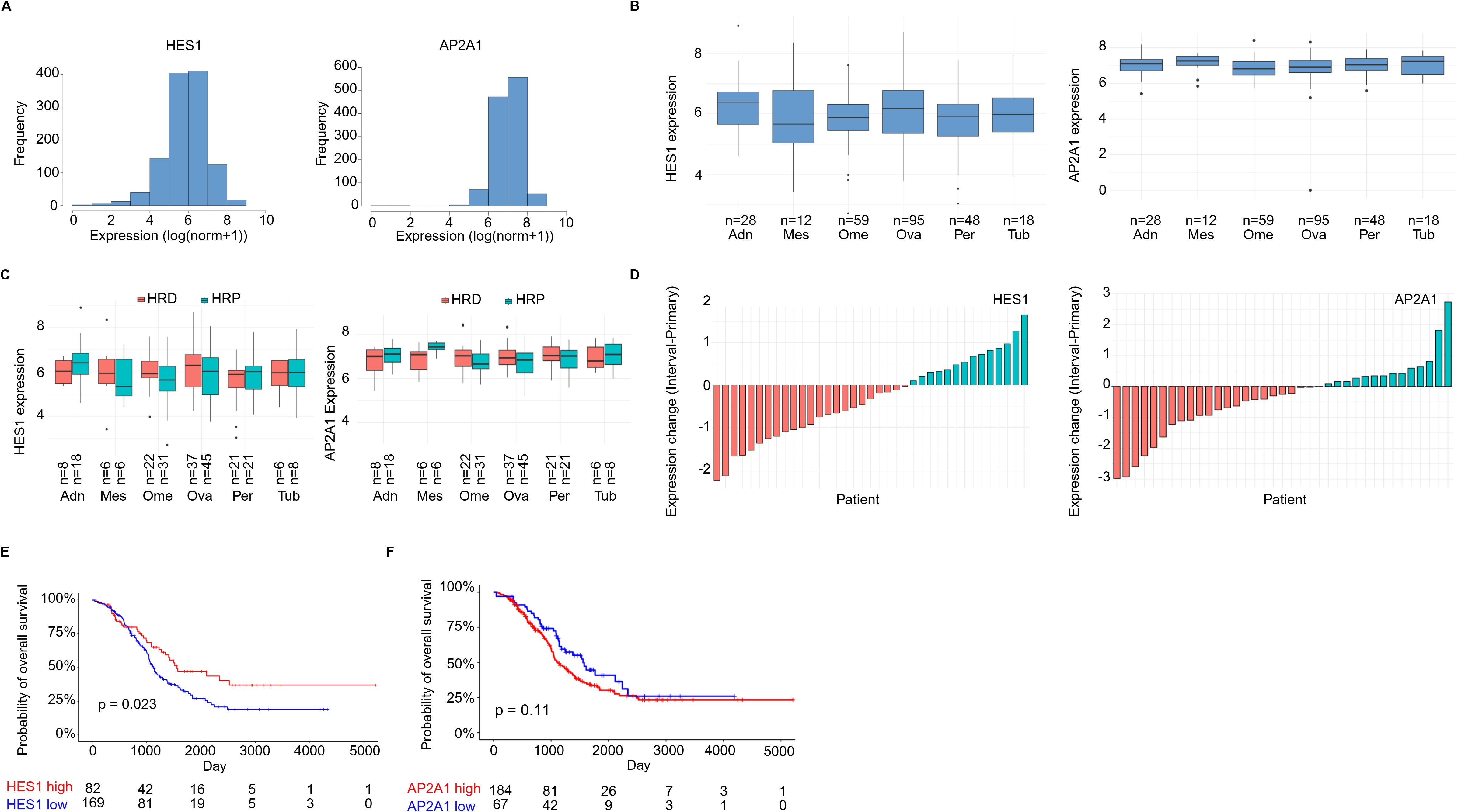
Clinical relevance of HES1 expression in HGSC. A Expression levels of HES1 and AP2A1 in the DECIDER HGSC cohort. B HES1 expression across tumor samples derived from different anatomical sites. Adn: adnexa; Mes: mesentery; Ome: omentum; Ova: ovary; Per: peritoneum; Tub: fallopian tube. C HES1 expression in samples classified as homologous recombination–proficient (HRP) or homologous recombination–deficient (HRD). D Changes of HES1 or AP2A1 expression following primary chemotherapy. E-F Survival analyses of patients based on HES1 (E) and AP2A1 (F) expressions in the DECIDER cohort.

